# Membrane localisation and checkpoint blockade enhance xenoantigen delivery to redirect pre-existing immunity against tumours

**DOI:** 10.64898/2026.03.01.708859

**Authors:** Sylvie Hauert, Fuxin Zhou, Justas Sidiskis, Aishwarya Saxena, Zoe Goldberger, Kevin Chang, Chiara Kling, Natalie Köhler, Stefan Fichtner-Feigl, Jeffrey A. Hubbell, Huda Jumaa, Priscilla S. Briquez

## Abstract

Cancer immunotherapies often rely on the recognition of tumour antigens, which strongly limits their efficacy upon heterogeneous antigen expression or downregulation. A strategy to overcome this limitation is to redirect pre-existing antiviral immunity against tumours through the delivery of xenoantigens. While many studies have addressed this by repurposing licensed vaccines, we here investigated the underlying mechanisms of immune redirection via the delivery of non-adjuvanted xenoantigen proteins, thereby avoiding confounding adjuvant- or pathogen-specific effects. Using B16F10 melanoma cells engineered to express the model antigen OVA, we found that tumour rejection in pre-immunised mice depends on the subcellular localisation of the xenoantigen, with membrane-bound antigens eliciting stronger rejection than dose-matched soluble cytoplasmic antigens. Enhanced rejection of membrane-bound OVA expressing tumours was associated with stronger CD4^+^ T cell responses. In addition, pre-immunisation also increased recruitment of inflammatory monocytes and macrophages at the tumour site. To translate this concept therapeutically, we developed a membrane-targeting OVA fusion protein which, upon intratumoural delivery, redirected pre-existing immunity and made tumours responsive to anti-PD-1 therapy. Importantly, these findings were further validated using the clinically relevant varicella zoster virus (VZV) glycoprotein E (gE) antigen and the licensed varicella vaccine Varivax. Our approach provides a mechanistic and translational perspective for treating poorly immunogenic tumours, leveraging widespread pathogen-specific immune memory in combination with anti-PD1 therapy in cancer patients.

## INTRODUCTION

Over the past decades, cancer immunotherapies have renewed hope for durable cancer cures, demonstrating that the immune system can effectively be harnessed against tumours. Currently, most immunotherapies rely on the direct or indirect recognition of tumour antigens. For example, targeted monoclonal antibodies, chimeric antigen receptor (CAR) T cells, bispecific T-cell engagers (BiTEs), and therapeutic cancer vaccines are all designed to directly recognize tumour antigens^1–5^. Others, such as immune checkpoint blockade (ICB; e.g., anti-PD-1), act by reactivating endogenous anti-tumour immune responses, still indirectly depending on the presence of tumour antigens. Indeed, the efficacy of anti-PD-1 therapy positively correlates with tumour mutational burden (TMB), which generates neoantigens, and tumour infiltration by T cells^6,7^.

A key limitation of antigen-centric cancer immunotherapies is that tumours can evade the immune system by downregulating the targeted antigens or through tumour heterogeneity, where some cancer cells lack the expression of the target. Both mechanisms compromise therapeutic efficacy and promote cancer relapse. A strategy to overcome this challenge is to actively deliver a chosen antigen to the cancer cells rather than to rely on endogenous tumour antigens. Importantly, delivering an antigen already recognised by the immune system can redirect pre-existing immunity, rapidly boosting tumour rejection through the reactivation of immune memory^8^.

Worldwide, populations have acquired long-term immunity against many pathogens through natural infection or vaccination, providing a rich repertoire of xenoantigens for immune redirection in cancer patients^9^. For instance, cytomegalovirus (CMV) infects a large fraction of the global population, with CMV-specific antibodies and circulatory T cells readily detected in 50-83% of people^10,11^. As another example, about 75% of children globally have received complete measles vaccination, which is expected to confer lifelong immunity^12,13^. Many other vaccines have also induced broad protective immunity across populations, including those against diphtheria-tetanus-pertussis toxins (DTP), hepatitis B, poliomyelitis, Bacillus Calmette-Guérin (BCG; tuberculosis), human papillomavirus (HPV; cervical cancer), varicella zoster virus (VZV; chickenpox/shingles) and more recently SARS-CoV-2 (COVID-19)^8^. In cancer patients, this widespread pathogen-specific immune memory offers an opportunity to repurpose approved vaccines as a mean to redirect pre-existing immunity against tumours.

In this study, we aim to clarify how intratumoural delivery of xenoantigen can optimally redirect pre-existing immunity for cancer immunotherapy, allowing us to isolate the effects of the xenoantigen. Building on our previous findings that membrane-bound antigens enhance tumour immunogenicity and responsiveness to ICB^25^, we hypothesize that targeting xenoantigens to the cancer cell membrane would strengthen immune redirection and improve tumour control. Using B16F10 melanoma cells engineered to express the model antigen OVA in membrane-bound or cytoplasmic soluble form, we first demonstrated that pre-immunised mice rejected tumours more effectively when OVA was membrane-bound, a difference driven by CD4^+^ T cells. Furthermore, we showed that pre-immunisation substantially increased the recruitment of inflammatory monocytes and macrophages at the tumour site. To translate this concept therapeutically, we designed an antibody fragment (Fab)-OVA fusion protein targeting the B16F10 cell surface. Although the fusion protein did not outperform OVA *in vivo*, intratumoural delivery of this xenoantigen sensitised B16F10 tumours to anti-PD-1, with enhanced effects in pre-immunised host. Finally, we confirmed the efficacy of this dual therapy – xenoantigen delivery combined with anti-PD1 – in pre-immunised hosts using the clinically relevant VZV glycoprotein E (gE) or via the licensed Varivax^®^ (VZV Oka strain) vaccine. Our study establishes that xenoantigen delivery can be enhanced through membrane targeting and combination with anti-PD1 therapy to achieve potent immune redirection and tumour control, providing a promising strategy for immunotherapy of poorly antigenic cancers.

## RESULTS

### Membrane-targeted expression of xenoantigen enhances pre-existing immune redirection against tumours

First, we questioned how the subcellular localisation of a xenoantigen in tumour cells affects tumour growth in pre-immunised mice, particularly when expressed in membrane-bound or cytoplasmic soluble form in B16F10 cells. As a xenoantigen, we used the common model antigen ovalbumin (OVA). Mice were pre-immunised using prime-boost vaccinations with OVA admixed with the Th1-biasing adjuvant CpG-B, administered two weeks apart on day 0 and 14 **(Fig. 1A)**. Th1-skewed immune responses are known to enhance cell-mediated immunity, which is more effective against tumour cell killing. Eighteen days after vaccination, we observed high levels of OVA-specific antibodies in the plasma, with titers around 5-6 **(Fig 1B)**. The IgG1/IgG2c ratio was lower than 1 (IgG1/IgG2c = 0.83), confirming a slightly Th1-biased immune response. In addition, we confirmed the presence of OVA-specific CD8^+^ T cells in the blood of the mice after 18 and 35 days **(Fig. 1C, Fig. S1A)**. CD8^+^ T cells are generally regarded as the most cytotoxic T cells for tumour killing.

**Figure 1.**
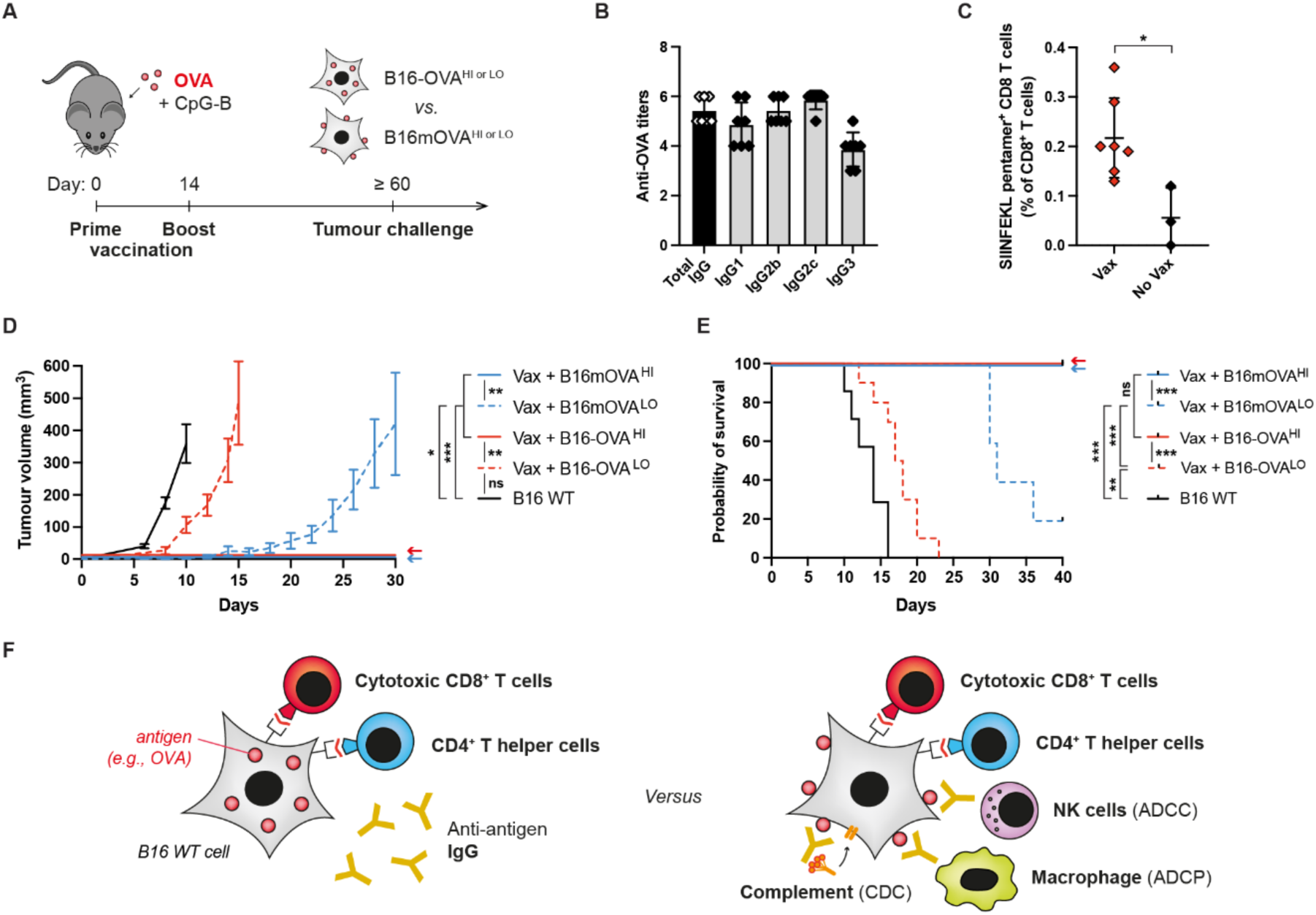
Membrane-targeted xenoantigen expression enhances tumour control in pre-immunised mice. **A)** Experimental timeline of the immunisation (i.e., vaccination) against OVA and subsequent tumour inoculation of the OVA-expressing B16F10 cell lines in mice. **B)** Titers of anti-OVA IgG per IgG subtypes in the plasma of the mice at d18. **C)** Percentage of OVA-specific CD8^+^ T cells against the immunogenic epitope OVA_257-264_ (SIINFEKL) in the blood at d18 in pre-immunised mice (vax) or naïve mice (no vax) (n ≥ 3, mean ± SD, unpaired t-test). **D, E)** B16F10 melanoma cells, either wild-type (WT) or genetically modified to express high (^HI^) or low (^LO^) doses of membrane-bound OVA (B16mOVA) or soluble OVA (B16-OVA), were injected intradermally in C57BL/6 mice. Tumour growth (D) and associated survival (E) of the different OVA-expressing B16 cell lines in pre-immunised mice (n ≥ 5, mean ± SEM, Kruskal-Wallis with Dunn’s post-test on d10, d14 or d26 for tumour growth, log-rank tests for survival). F) Potential anti-tumour mechanisms engaged by soluble (left) or membrane-bound (right) xenoantigens when expressed by cancer cells.

Following pre-immunisation, mice underwent a rest period of at least 46 days without OVA exposure to allow contraction of the acute anti-OVA response, leaving predominantly a memory immune response^26^. After this, mice were injected intradermally with B16F10 melanoma cells engineered to stably express OVA in an exoleaflet membrane-bound (B16mOVA) or a cytoplasmic soluble form (B16-OVA), at matching high (^HI^) or low (^LO^) expression levels, as shown by qPCR and flow cytometry **(Fig. S1B, C)**. Strikingly, we observed that all tumours expressing high doses of OVA were rejected, while tumours with low doses of antigens grew, although slower than the parental B16F10 wild-type (WT) cells **(Fig. 1D)**. Furthermore, we found that tumours expressing low doses of membrane-bound OVA grew with a significant >10 days delay as compared to the one with soluble OVA, supporting our hypothesis that membrane-targeted antigens in cancer cells enhance tumour immunogenicity and effectively redirect pre-existing immunity. The survival of these mice was consistent extended: 100% of the mice that received B16F10 tumours expressing high doses of OVA were cured **(Fig. 1E)**. Mice injected with B16mOVA^LO^ tumours showed a significantly extended survival of about 13 days (73% extension) as compared to mice that received B16-OVA^LO^ tumours, and of 16 days (114% extension) as compared to B16 WT injected mice. As a note, we previously published that all these cell lines were tumorigenic and lethal in naïve (i.e., non-pre-immunized) mice, supporting that tumour rejections are mediated by the host pre-existing immunity **(Fig. S1D)**.

They are several reasons that could explain why membrane-targeting of the xenoantigen could improve tumour rejection as compared to soluble expression of xenoantigen. First, the antibodies present in high levels in the plasma after mice pre-immunisation may bind to the membrane-bound xenoantigen and support multiple anti-tumour immune mechanisms. Indeed, natural killer (NK) cells may enhance the release of cytotoxic granules via antibody-dependent cell cytotoxicity (ADCC), macrophages may boost their phagocytosis via antibody-dependent cell phagocytosis (ADCP), and the antibodies themselves could induce tumour cells lysis via the activation of the complement system (CDC)^1^. Importantly, the endogenous production of the xenoantigen by the tumour cells may lead to its direct presentation on the major histocompatibility complex (MHC)-I and -II (via autophagy) at the surface of the tumour cells, allowing their direct recognition by CD8^+^ and CD4^+^ T cells, respectively **(Fig. 1F)**. As a note, although not all tumours express MHC-II, we have previously shown the parental B16F10 cells and the engineered OVA-expressing B16F10 cell lines express MHC-I and MHC-II at similar level^25^.

### Superior tumour rejection by membrane-targeted xenoantigen expression depends on CD4^+^ T cells

To understand the underlying immune mechanisms by which membrane-targeted xenoantigens improved tumour rejection in pre-immunised hosts, we performed a set of functional immune depletion experiments *in vivo*. We started by evaluating rejection of the most immunogenic B16mOVA^HI^ tumour when implanted in pre-immunised nude mice, lacking mature T cells, and observed that the tumours developed similar to the B16F10 WT tumours **(Fig. 2A)**. As expected, nude mice were uncapable to generate anti-OVA IgG during the pre-immunisation **(Fig. S2A)**. This indicates that full rejection of B16mOVA^HI^ in the pre-immunised WT mice observed in Fig. 1D relies on the immune redirection of OVA-specific T cell responses.

**Figure 2.**
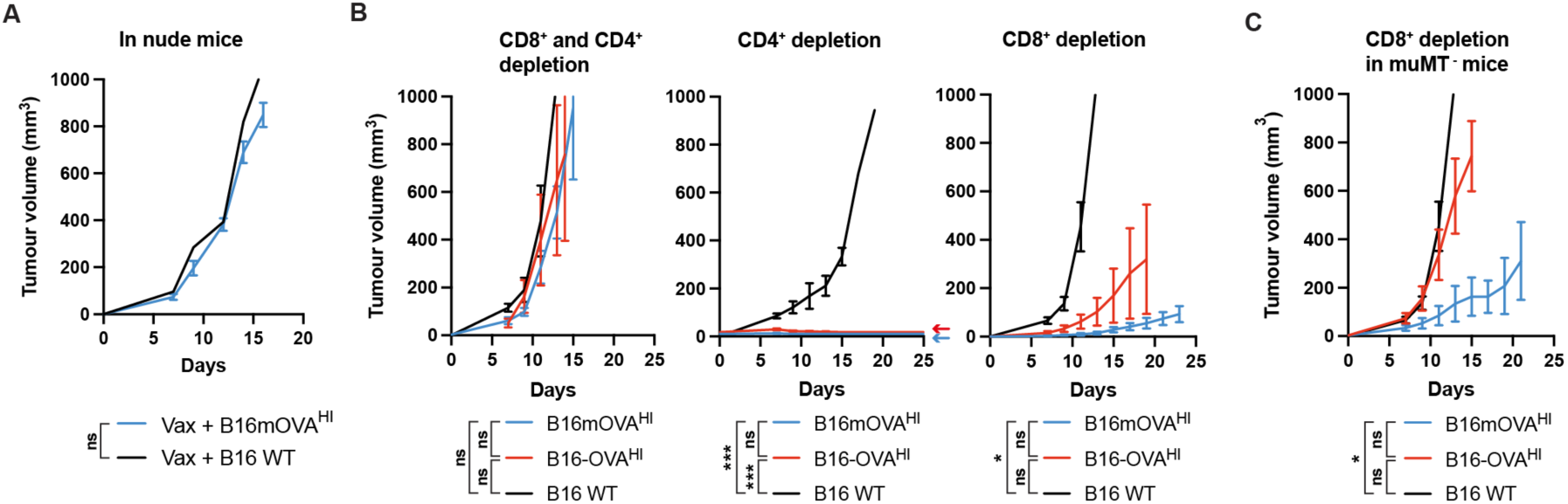
Membrane-targeted xenoantigens improve anti-tumour CD4^+^ T cell responses and potentially enhance antibody-mediated cytotoxic mechanisms in pre-immunised mice. **A)** Tumour growth of B16mOVA^HI^ in nude mice, lacking mature T cells (n ≥ 2, mean± SEM, Kruskal-Wallis with Dunn’s post-test on d14). **B)** Tumour growth of B16mOVA^HI^ in mice depleted of CD8^+^ and/or CD4^+^ cells (n ≥ 3, mean ± SEM, Kruskal-Wallis with Dunn’s post-test on d15). **C)** Tumour growth of B16mOVA^HI^ in muMT^-^ mice depleted of CD8^+^ cells (n ≥ 4, mean ± SEM, Kruskal-Wallis with Dunn’s post-test on d15).

To isolate the effect of CD8^+^ and CD4^+^ T cells, we then performed experimentation in WT mice using depleting anti-CD8 and anti-CD4 antibodies. Pre-immunised mice were treated with anti-CD8, anti-CD4 or both depleting antibodies 3 days before tumour inoculation, and again at day 4 and 11 post-inoculation. We then validated that CD8^+^ and/or CD4^+^ T cells were successfully depleted of the mice blood after treatment **(Fig. S2B)**. We observed that B16mOVA^HI^ and B16-OVA^HI^ grew at similar rate as the B16F10 WT tumours in mice depleted of both CD8^+^ and CD4^+^ T cells **(Fig. 2B)**, consistent with our observation in the nude mice. We then showed that pre-immunised mice depleted of CD4^+^ T cells were capable of fully rejecting both the B16mOVA^HI^ and B16-OVA^HI^ tumours **(Fig. 2B)**, like in the WT mice **(Fig. 1D)**. In contrast, pre-immunised mice depleted of CD8^+^ T cells were incapable of eradicating B16mOVA^HI^ and B16-OVA^HI^ tumours. In these mice, B16mOVA^HI^ tumours grew significantly slower than the B16F10 WT ones, whereas B16-OVA^HI^ did not **(Fig. 2B)**. These results reveal that although the CD8^+^ T cells are the dominant cytotoxic effectors redirected against tumours, CD4^+^ T cells can also substantially control tumour growth, particularly when the xenoantigen is membrane-bound. This further suggests that the superior rejection of tumours bearing membrane versus soluble ones – especially at low doses of OVA expression **(Fig. 1D, E)** – are driven by differences in CD4+ T cell responses.

Lastly, to determine whether antibodies play a role in the enhanced rejection of the B16mOVA^HI^, we repeated the CD8^+^ T cell depletion experiment in muMT^-^ mice. These mice constitutively lack mature B cells and were therefore uncapable to generate antigen-specific IgG during the pre-immunisation **(Fig. S2A)**. Again, we observed that the growth of B16mOVA^HI^ tumours, but not of B16-OVA^HI^, was significantly reduced compared to B16F10 WT tumours **(Fig. 2D)**. This confirms that the enhanced rejection of B16mOVA^HI^ tumours depends on CD4+ T cells, independently of the presence of antigen-specific antibodies. Nevertheless, we noticed that the growth of B16mOVA^HI^ and B16-OVA^HI^ was accelerated in muMT^-^ mice compared to WT mice **(Fig. 2B, C)**, suggesting that antibodies contribute to tumour control, although they do not appear to be a major mechanism of anti-tumour immunity at play **(Fig. S2C)**.

### Pre-existing immunity enhances recruitment of inflammatory monocytes and macrophages in membrane-targeted xenoantigen-expressing tumours

We then asked whether innate immunity was also modulated by pre-immunisation in mice and how could this contribute to enhanced tumour rejection. To study this, the skin of mice injected with B16mOVA^HI^ cells was collected 4 days post-inoculation, and the innate cell populations were analysed by flow cytometry **(Fig. S3)**. We compared the injected site of mice that were pre-immunized against OVA, naïve (i.e., non-pre-immunised), transgenic Act-mOVA (i.e., immune tolerant to OVA), and non-injected with tumour cells (i.e., no tumour, healthy skin only). We first showed that injection of tumour cells increased the recruitment of innate CD11b^+^ myeloid cells as compared to healthy skin, independently of the pre-immunisation **(Fig. 3A)**. Then, we observed that pre-immunisation significantly enhanced infiltration of inflammatory monocytes and macrophages, particularly of the MHC-II^+^ macrophages, which are expected to be more activated/pro-inflammatory and better armed to act as antigen-presenting cells (APCs) for the CD4^+^ T cells **(Fig. 3B)**. In addition, myeloid cells expressing FcγRII and FcγRIII were also more numerous in the tumours in pre-immunised mice. These cells can bind the Fc domain of antibodies and possibly use them to perform antibody-dependent anti-tumour mechanisms (e.g., ADCP) and mostly include monocytes-derived macrophages in this context **(Fig. 3C)**. Indeed, we did not find any differences in infiltrating neutrophils, dendritic cells (DCs) or NK cells as a result of pre-immunisation. Together, we found that host pre-immunisation increased innate inflammation in the microenvironment of tumours expressing the xenoantigen, as shown here using the membrane-targeted antigen expression model.

**Figure 3.**
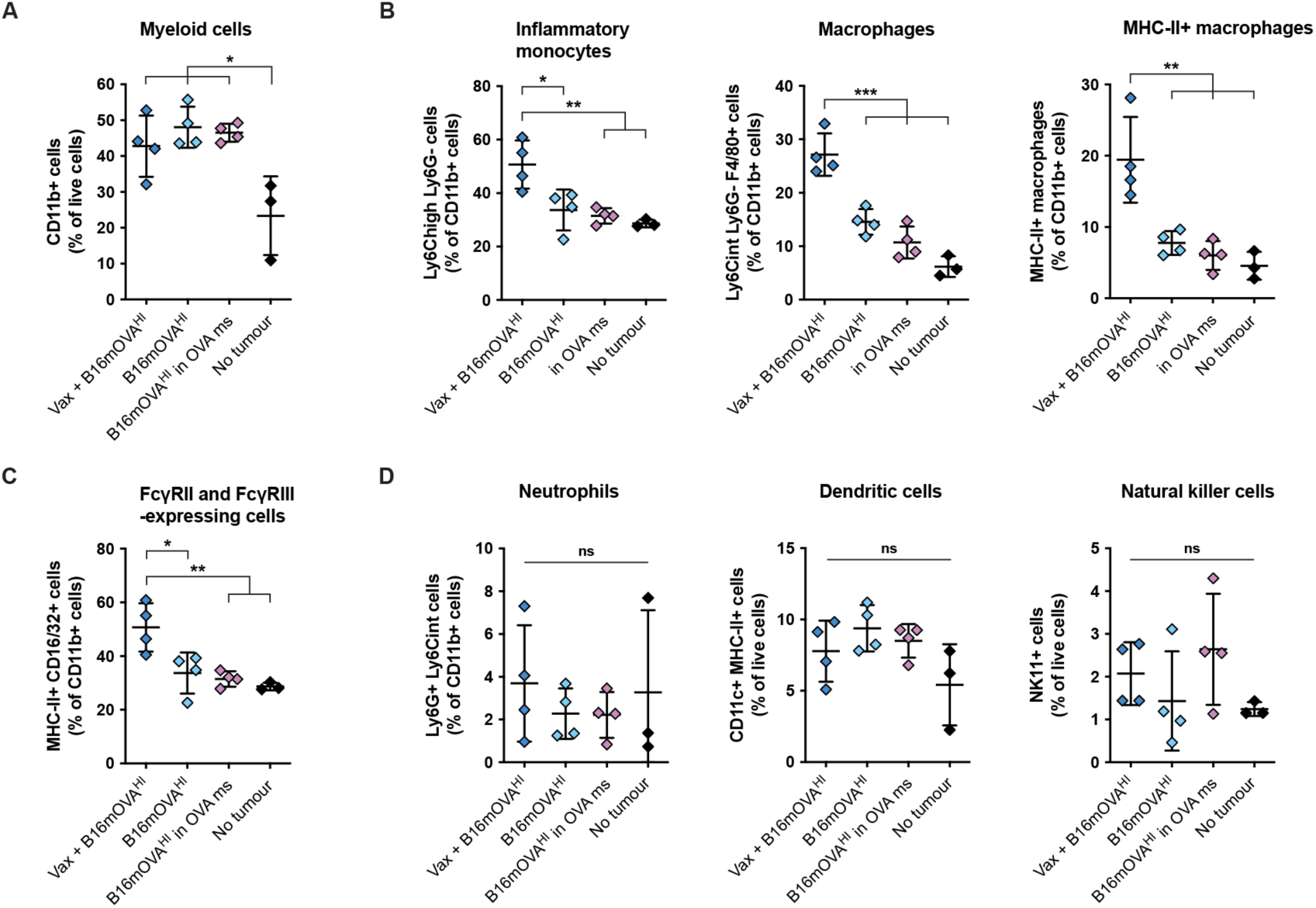
B16mOVA^HI^ tumours increase infiltration of inflammatory monocytes and macrophages at the tumour site. Innate immune cells were quantified by flow cytometry in the skin of pre-immunised (Vax), naïve or Act-mOVA mice (OVA ms), at the site of B16mOVA^HI^ inoculation, 4 days after tumour injection. **A)** Quantification of myeloid cells (n ≥ 3, mean ± SD, ANOVA with Tukey’s post-test). **B)** Quantification of inflammatory monocytes and macrophages (n ≥ 3, mean ± SD, ANOVA with Tukey’s post-test). **C)** Quantification of immune cells expressing FcγRII/III (CD16/32^+^) at the tumour site (n ≥ 3, mean ± SD, ANOVA with Tukey’s post-test). **D)** Quantification of neutrophils, dendritic cells and natural killer cells.

### Intratumoural delivery of recall xenoantigen combined with anti-PD-1 therapy enhanced tumour control

Next, we aimed to translate our findings into a therapeutic approach. Since we showed benefits of membrane-targeted expression of xenoantigen on tumour control **(Fig. 1D, E, Fig. 2B, C)**, we designed a membrane-targeting protein-based approach to deliver at the surface of the B16F10 WT cells. We used an antibody against tyrosinase-related protein 1 (TRP1), taken from the clone TA99, which is a murine IgG2a antibody that enables ADCC and promotes tumour clearance upon membrane targeting of melanoma^27–29^. We first confirmed the effective binding of TA99 antibody to B16F10 tumours by immunohistochemistry **(Fig. 4A)** and validated *in vivo* targeting in the B16F10 WT tumours through its ability to delay tumour growth **(Fig. 4B)**. Then, we used TA99’s variable regions to generate a Fab fragment and genetically fused it to the OVA full-length protein **(Fig. 4B)**. This protein, called FabTRP-OVA, was produced in human embryonic kidney (HEK)-293 cells and successfully purified by affinity chromatography toward histidine-tag and the Fab constant regions **(Fig. 4C)**. We additionally demonstrated by flow cytometry that FabTRP-OVA effectively bound to the surface of the B16F10 WT cells, in contrast to wild-type OVA **(Fig. 4D, E)**.

**Figure 4.**
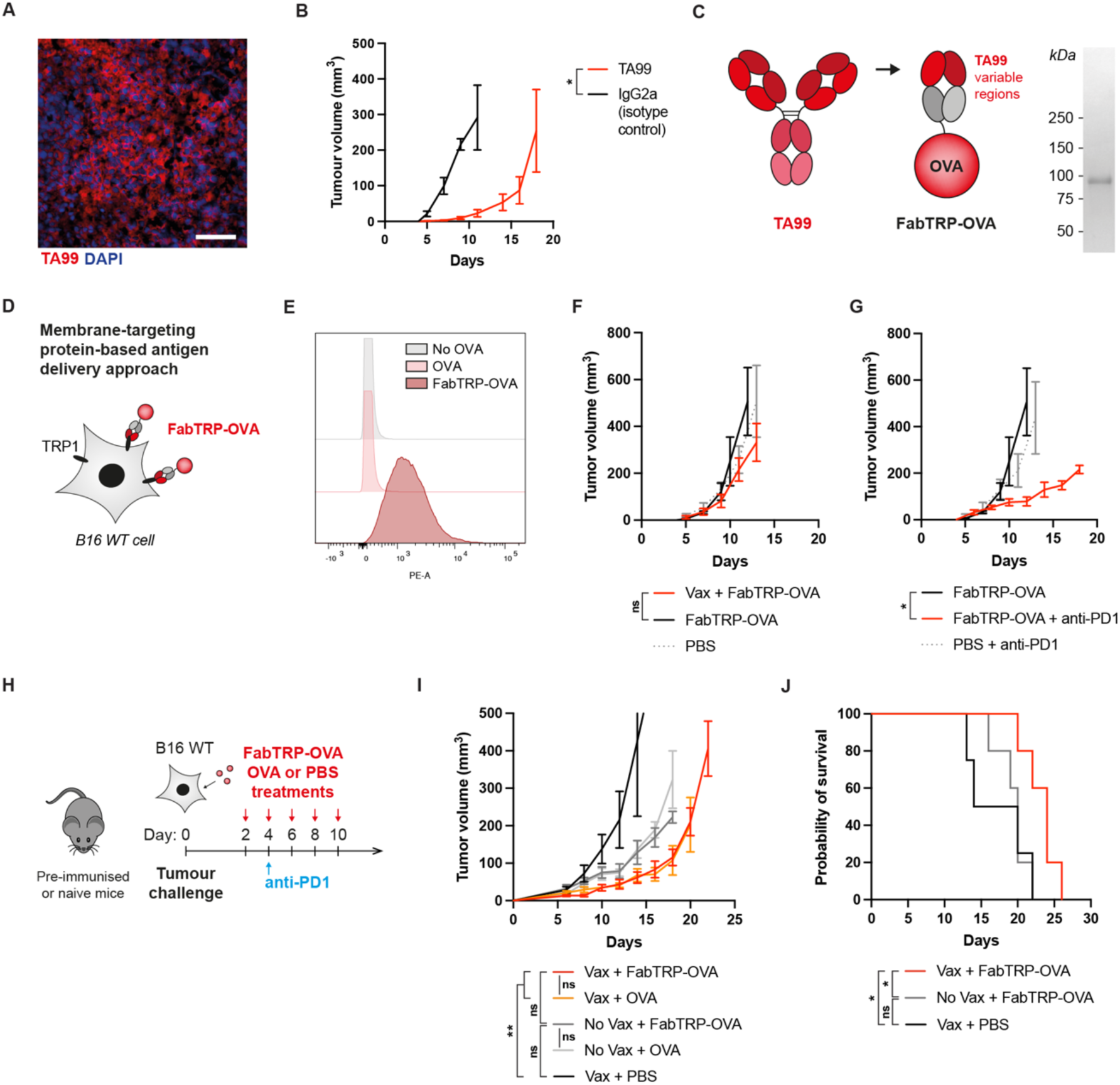
Delivery of OVA xenoantigens in pre-immunised tumour-bearing mice, exploring membrane-targeting protein-based delivery approach. **A)** Immunofluorescent staining of B16F10 WT melanoma with the TRP1-targeting TA99 antibody (scale bar = 100 μm). **B)** B16F10 WT tumour growth upon treatment with TA99 or IgG2a isotype control (n ≥ 4, mean ± SEM, Mann-Whitney test on d11). **C)** Design of FabTRP-OVA fusion protein and analysis of its production by SDS-PAGE (expected size = 92,3 kDa). **D)** Illustration of the FabTRP-OVA fusion protein approach for targeting the xenoantigen to the cancer cell membrane. **E)** Representative flow cytometry plot of FabTRP-OVA, OVA and no OVA (i.e., secondary antibody only) binding to the surface of B16F10 WT cells. **F)** B16F10 WT tumour growth when treated with FabTRP-OVA in pre-immunised or naïve mice (n ≥ 4, mean ± SEM, Kruskal-Wallis with Dunn’s post-test on d12). **G)** B16F10 WT tumour growth when treated with FabTRP-OVA in combination with anti-PD-1 checkpoint blocade therapy (n ≥ 4, mean ± SEM, Kruskal-Wallis with Dunn’s post-test on d12). **H)** Experimental treatment timeline of xenoantigen delivery (i.e., FabTRP-OVA, OVA or PBS (vehicle only)) and anti-PD-1 checkpoint blockade therapy in B16F10 WT in mice pre-immunised against OVA (Vax) or naïve (No vax). **I, J)** B16F10 WT tumour growth (I) and associated mouse survival (J) upon treatment with FabTRP-OVA, OVA or PBS in combination with anti-PD-1 checkpoint blockade therapy (n ≥ 5, mean ± SEM, Kruskal-Wallis with Dunn’s post-test on d16, log-rank tests for survival).

We then assessed the therapeutic efficacy of FabTRP-OVA when delivered intratumourally in B16F10 WT tumours. FabTRP-OVA was given at early stage of tumour development, starting at day 2 post-inoculation and repeated every 2 days for a total of 5 doses. We observed that FabTRP-OVA has no therapeutic efficacy as a stand-alone treatment on tumour growth in naïve mice, nor in pre-immunised mice **(Fig. 4F)**. We reasoned that the xenoantigen was here delivered after B16F10 WT inoculation, so potentially in an already immunosuppressive microenvironment, which could hinder the immune-related effects of FabTRP-OVA. This contrasted with our previous model, in which the tumours constitutively expressed the OVA at the time of the tumour inoculation. Since tumour rejection requires the effective T cell responses **(Fig. 2A-C)**, we next decided to combine FabTRP-OVA treatment with anti-PD-1 therapy.

After confirming that the B16F10 WT tumours were unresponsive to anti-PD-1 as a stand-alone treatment **(Fig. S4A)**, we treated naïve mice with intratumoural delivery of FabTRP-OVA and intraperitoneal administration of anti-PD-1. Strikingly, we observed that the combined therapy significantly slowed tumour growth **(Fig. 4G)**, showing that the xenoantigen sensitised B16F10 melanoma to anti-PD-1. We therefore assessed the effect of host pre-immunisation and of membrane-targeting in mice treated with anti-PD-1 **(Fig. 4H)**. We first observed that mice pre-immunisation combined with intratumoural xenoantigen delivery slowed tumour growth, regardless of the membrane-targeting, and extended mice survival as compared to both naïve mice treated with the xenoantigen and pre-immunised mice treated with PBS **(Fig. 4I, J**, **Fig. S4B)**. Second, we found that targeting OVA to the cancer cell membrane using our protein fusion approach did not provide benefit as compared to the delivery of wild-type OVA, independently of the mice pre-immunisation status **(Fig. 4I)**. This highlighted that the fusion protein-based targeting approach we explored was not effective in these experimental conditions and could not recapitulate the benefits observed upon constitutive expression of membrane-bound antigen **(Fig. 1D)**. While there are many parameters that could be optimized to improve membrane-targeting strategy, which are later addressed in the Discussion, we chose at this point to further focus on the characterization of the combination therapy of xenoantigen delivery and anti-PD-1 in pre-immunised mice, which successfully enhanced tumour control. This permitted the use of licensed vaccines, without the need for engineering of the vaccinal antigen.

### Delivery of the relevant *VZV* antigen gE or vaccine Varivax with anti-PD-1 therapy enhances tumour control in pre-immunised mice

We next asked whether the benefits of combining xenoantigen delivery with anti-PD-1 applied to other antigens than OVA. To test this, we selected a VZV antigen from the Oka strain (VZVO) – the causative agent of the chickenpox – against which many individuals are immunised through infection or vaccination. Globally, about 4.5 million cases of chickenpox were reported among older adults in 2021^30^, and in the USA, vaccination campaigns have prevented an estimated about 91 million cases (97%)^31^. Among viral proteins, the glycoprotein E (gE) is the one known to mediate virus entry into human cells, and it is one of the most abundant and immunodominant antigen^32^.

To pre-immunise mice, we used the Varivax^®^ vaccine (Merck), a live-attenuated VZV Oka strain, in which viral glycoproteins are incorporated into the virus envelope, and which contains natural immune-stimulating viral components that acts as adjuvants. Mice were vaccinated with 13.5 plaque-forming units (pfu) of Varivax at day 0 and 14, or with an in-house-made vaccine containing 12 μg of gE and 25 μg of CpG-B (similar to the previously used OVA vaccine) **(Fig. 5A)**. We found that mice generated IgG at similar levels with both formulations, analysed in the plasma at day 55 **(Fig. 5B)**. We then evaluated cellular responses to these vaccines. To do so, mice received an additional boost vaccine using gE/CpG-B at day 60 and their spleen was collected 5 days later. Splenocytes were restimulated *in vitro* with gE and the amount of gE-specific T cells was determined via the secretion of IFNγ. We observed that both Varivax and the gE/CpG-B vaccinations were able to generate gE-specific T cells to similar levels, in contrast to naïve mice **(Fig. 5C)**. As a note, it is known that VZV is highly specific to human cells and does not infect mouse cells; therefore, the observed T cell response is likely to be biased toward CD4^+^ T cells^33^.

**Figure 5.**
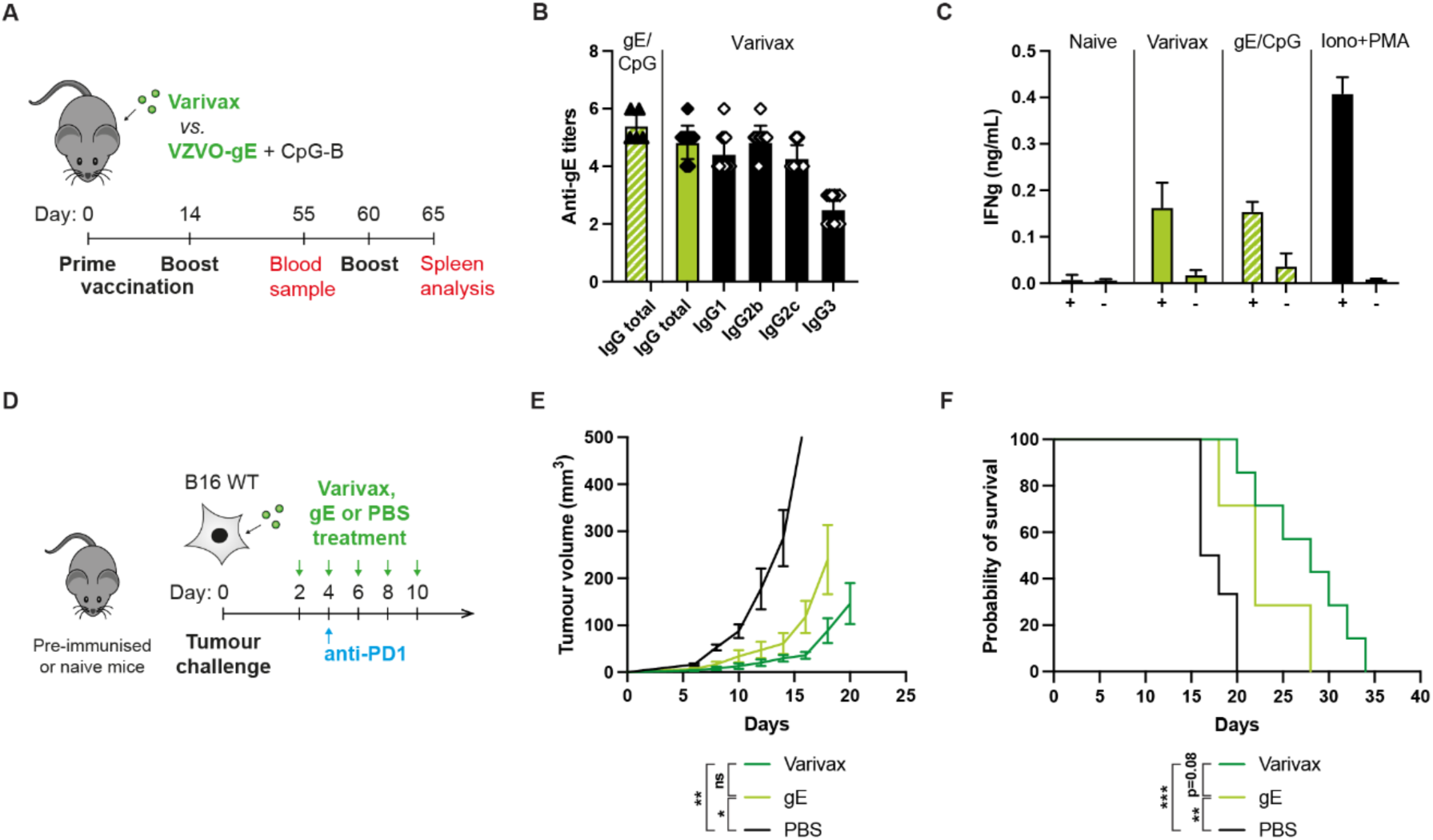
Delivery of VZVO-gE xenoantigens in pre-immunised tumour-bearing mice. **A)** Experimental timeline of the immunisation (i.e., vaccination) against *varicella VZVO* using Varivax or a combination of gE/CpG. **B)** IFΝγ quantification in the supernatant of gE restimulated (+) or non-restimulated (-) splenocytes, collected from mice immunized with Varivax, gE/CpG or PBS (i.e., naïve). Ionomycin+PMA was used as a positive control. **C)** Titers of anti-gE total IgG and per IgG subtypes in the plasma of the pre-immunised mice at d55 (before tumour implantation). **D)** Experimental treatment timeline of xenoantigen delivery (i.e., Varivax, gE or PBS (vehicle only)) and anti-PD-1 checkpoint blockade treatment in B16F10 WT in mice pre-immunised with Varivax against *varicella VZVO*. **E, F)** B16F10 WT tumour growth (E) and associated mouse survival (F) upon treatment with Varivax, gE or PBS in combination with anti-PD-1 checkpoint blockade (n ≥ 7, mean ± SEM, Kruskal-Wallis with Dunn’s post-test on d16, log-rank tests for survival).

Next, we tested the therapeutic efficacy of intratumoural gE or Varivax delivery in mice pre-immunised with Varivax and treated with anti-PD-1 **(Fig. 5D)**. Consistent with our previous results using OVA, mice treated with non-adjuvanted gE or the Varivax vaccine exhibited significantly slower tumour growth compared to PBS-treated controls **(Fig. 5E)**, which led to extended survival **(Fig. 5F)**.

We also observed a non-significant trend toward greater efficacy of Varivax compared recombinant gE. There are many possible reasons for this trend, particularly that 1) the gE dose in the Varivax group differs from the one in the recombinant gE group, 2) Varivax contains innate immune-stimulatory components from the live attenuated virus, whereas the recombinant gE was non-adjuvanted, 3) Varivax contains multiple antigens, not only gE, and 4) the antigens in Varivax are membrane-associated, in contrast to the soluble recombinant gE.

Overall, these results demonstrate that intratumoural vaccination against an immune recall xenoantigen – here exemplified using the clinically relevant Varivax vaccine – effectively controls tumours when combined with anti-PD-1 therapy.

## DISCUSSION

Tumours with poor antigenicity represent a challenge for cancer immunotherapies, which have generally been designed to target specific tumour antigens. In this study, we querried whether in such tumours, the delivery of recombinant xenoantigens would effectively redirect pre-existing immunity and enhance tumour control. We selected the aggressive and poorly immunogenic B16F10 melanoma as a study model. In our previous work, we found that the subcellular localisation of tumour antigens was a key parameter to determine tumour immunogenicity^25^. Indeed, tumours expressing membrane-bound OVA were enriched in immune cells compared to those expressing soluble OVA, particularly in CD8^+^ T cells and NK cells, but not in CD4^+^ T cells or macrophages. In addition, they secreted higher levels of IFNγ, TNFα and other NK/T cells recruiting chemokines (e.g., CCL-3, CCL-4, CXCL-9, CXCL-10), creating a more pro-inflammatory tumour microenvironment.

Here, we demonstrated that in pre-immunised mice, B16F10 cells expressing high levels of xenoantigen (i.e., OVA) were fully rejected, regardless of antigen localisation, whereas at lower expression levels, membrane-bound OVA enabled markedly better tumour rejection. Because these tumours all grew in naïve mice, these results highlight that pre-existing immunity against OVA was effectively redirected against the tumour cells in pre-immunised hosts. Mechanistically, we found that tumours were primarily rejected by CD8^+^ T cells, irrespective of the antigen localisation. In addition, we highlighted that tumours expressing membrane-bound antigen elicited a stronger CD4^+^ T cell response, which could explain the superior rejection of these tumours at low antigen expression levels. It has been showed that immune redirection in pre-immunised host generally relies on T cell responses^11,17–23^. Interestingly, a few studies repurposing vaccines against tumour have specifically underscored a pivotal role for CD4^+^ T cells, not only supporting antiviral CD8^+^ T cells, but also in directly boosting tumour-specific CD8^+^ T cell cytotoxicity^21,24,34–36^. In addition, we observed that pre-existing immunity increased the recruitment of inflammatory monocytes and macrophages at the tumour-injected site, a finding in line with other models where recall responses boosted monocyte recruitment and macrophage activity via CCL-3 secretion by memory CD8^+^ T cells^37^. As previously mentioned, B16F10 tumours expressing membrane-bound antigens were also particularly enriched in CD8^+^ T cells, with elevated CCL-3 levels^25^.

Taken together, a possible explanation for the enhanced rejection of the membrane-bound OVA tumours in pre-immunised mice is that the stronger activity of OVA-specific Th1 CD4^+^ T cells within the tumour, mediated by the detection of OVA epitopes presented on the MHC-II of the B16F10 themselves or of other APCs, leads to increased secretion of pro-inflammatory cytokines^25^. These cytokines may in turn enhance MHC molecule expression and OVA epitope presentation, as well as enhance local inflammation in the tumour. This increases the recruitment of inflammatory monocytes and macrophages and enhances anti-tumour activity, a process that could further supported by the pre-existing OVA-specific antibodies, in case of ADCP^38^. CD4^+^ T cells may enhance both OVA- and tumour-specific CD8^+^ T cells, and could exert direct cytotoxicity, although the latter is generally limited in mouse models^39,40^. Moreover, other studies have shown that membrane-bound or particulate antigens further enhance B cells responses and cross-presentation by DCs compared to soluble antigen^41–43^.

Although we used OVA as a model xenoantigen, our findings based on the OVA-expressing B16F10 cell lines can also be interpreted in the context of tumour neoantigens. Since pre-existing immunity was far more effective against tumours expressing membrane-bound OVA, our data suggest that preventive vaccination targeting membrane-bound tumour neoantigens could be substantially more effective than vaccines against soluble cytoplasmic neoantigens. Currently, strategies for preventive cancer vaccine development select antigens that are known to be highly expressed in tumours, without much consideration of their subcellular localisation^44^. We believe that incorporating this criterion could markedly enhance preventive cancer vaccine efficacy.

We then sought to translate the benefits of membrane-targeted xenoantigen expression into a therapeutic strategy. To do so, we explored the intratumoural delivery of a membrane-targeting OVA fusion protein (i.e., FabTRP-OVA), which successfully bound to B16F10 cells, yet did not confer therapeutic benefits over the WT OVA *in vivo*. This lack of difference may reflect the experimental dosing schedule, as overly frequent intratumoural injections could saturate the tumour with the antigen, masking the benefits of membrane targeting. Furthermore, the protein fusion approach may fail to replicate the effects seen upon cellular expression of the antigen, as endocytosis and presentation of OVA epitopes on MHC molecules are less efficient with protein delivery^45^. Lastly, anti-OVA pre-existing neutralizing antibodies might have prevent binding of FabTRP-OVA to the membrane *in vivo*^46,47^. That said, multiple strategies exist to induce xenoantigen expression in cancer cells, including viral vectors (e.g., live-attenuated viruses, adenoviruses, adeno-associated virus, nanoparticles (e.g., lipid-based nanoparticles LNPs), gene electroporation, plasmid-based gene delivery. Determining the most appropriate approach for membrane localised expression of xenoantigen would require systematic assessment and direct comparison, which is beyond the scope of this study. Nevertheless, our study and other evidence suggests that platforms fostering strong CD4^+^ T cell responses may hold superior therapeutic potential for this purpose^18,21,23,48,49^.

Another potentially powerful approach to enhance intratumoral xenoantigen therapy in the context of pre-existing immunity is to combine it with anti-PD-1 therapy. In established tumours, the immunosuppressive microenvironment limits T cell activities, which can be reactivated by ICB. In our study, xenoantigen delivery in pre-immunised mice did not provide therapeutic benefits in the absence of anti-PD-1. Previous reports have already shown benefits of combining ICB with intratumoural vaccination using licensed vaccines^17,20,24,50,51^, Indeed, intratumoural vaccination has frequently been shown to convert poorly antigenic, non-inflamed “cold” tumours into inflamed “hot” tumours responsive to ICB^8,16,17^. However, we here highlight that a single, non-adjuvanted recombinant xenoantigen is sufficient to achieve this effect, in absence of adjuvants and pathogen-derived components present in vaccine formulations. Clinically, our data provides a rational for dual therapy of recall immunogenic xenoantigen delivery and anti-PD-1, considering that cancer patients already harbour pre-existing immunity against known pathogens, which could be redirected^52^. Recently, an alternative strategy has been proposed to redirect pre-existing immunity against tumours via the delivery non-adjuvanted viral epitope peptides for direct presentation on MHC molecules. This strategy has been successfully validated using viral peptides from CMV and Epstein-Barr virus (EBV), both widespread in populations, even in absence of ICB treatment^11,22,35,48,53–56^. Nevertheless, Rosato et al. also pointed that combining such viral epitope delivery with ICB further enhance tumour rejection^20^, reinforcing the rationale for the dual therapy. However, unlike viral epitope peptides that bind only to specific MHC molecules, whole antigenic proteins can be processed and presented across patients, increasing to the versatility of our approach.

Finally, we validated our findings using the clinically relevant VZVO gE antigen or the licensed Varivax vaccine. To our knowledge, repurposing Varivax for immune redirection in cancer therapy had not yet been investigated, although the VZV-vaccine Shingrix^®^ had recently shown to be effective in the TC-1 lung epithelial cancer in mice^23^. This suggests that our observations in the B16F10 melanoma model may be generalizable to other cancers.

As a conclusion, this study highlighted that membrane-targeting of xenoantigens on cancer cells and combination with ICB therapy enhanced intratumoural xenoantigen delivery and improve the redirection of pre-existing immunity, paving the way to effective immunotherapy against poorly immunogenic tumours.

## ACKNOWLEDGEMENTS

The authors would like to thank Prof. Melody A. Swartz for her scientific advice, and Susana Gomez, Yue Wang, Trevin Kurtanich and Grégoire Repond for their technical help. In addition, we acknowledge the Lighthouse Core Facility, partly funded by the Medical Faculty, University of Freiburg.

## AUTHOR CONTRIBUTIONS

S.H., F.Z., Z.G., K.C., J.A.H., and P.S.B., have conceived the project and designed the experiments; F.Z., J.S., A.S., C.K., H.J. performed *in vitro* experimentation; S.H. and P.S.B. performed *in vivo* experimentation; S.H., F.Z., N.K., S.F-F. and P.S.B. analysed and interpreted the data; Z.G., H.J. and P.S.B. wrote the manuscript, and K.C., N.K., S.F-F. and J.A.H proof-read it.

## MATERIALS AND METHODS

### Ethic statement

All animal experimentation were performed in accordance with the procedures approved by the University of Chicago Institutional Animal Care and Use Committee (IACUC) at the University of Chicago (IL, USA) in compliance with local ethical and procedural regulations, under the protocol n°72456.

### Mice

Female mice from the strains C57BL/6J (wild-type WT; No 000664), Nu/J (Nude; No 002019) B6.129S2-Ighm^tm1Cgn^/J (MuMt^-^; No 002288), and male mice C57BL/6J-Tg(CAG-OVAL)916Jen/J (Act-mOVA; No 005145) were obtained from The Jackson Laboratory (Bar Harbor, ME, USA). At the start of the experiment, all mice were between 8–12-week-old except the Act-mOVA that were around 30-week-old.

### Mice immunisation

Mice were anesthetised by isoflurane inhalation. For immunization against OVA, mice were injected with 10 μg of endotoxin-free OVA (EndoGrade, Hyglos, Regensburg, Germany) admixed with 25 μg of CpG-B (Trilink, BioTechnologies, San Diego, CA, USA) in sterile phosphate-buffered saline (PBS) in the 2 front hocks. For immunization against *varicella* VZVO, mice were injected with 13.5 pfu of the Varivax^®^ vaccine (Merck & co, Rahway, NJ, USA) diluted in sterile PBS. For immunization against the glycoprotein E (VZVO-gE), mice were injected with 12 μg recombinant VZVO-gE admixed with 25 μg of CpG-B. The prime vaccination (i.e., first injection) was administered at day 0 (d0) and defines the start of the experiment. A boost vaccination with the same formulation as the prime was then given on d14. No further treatment was administered until at least d60. Mice immunisation was performed in WT, nude and muMT^-^ mice, depending on the experiment.

### Detection of OVA-specific CD8^+^ T cells in the blood

At day 18 post-immunisation, 50 μL blood was taken from the mice via the saphenous vein and collected in an EDTA-treated tube. The blood was spined down at 400 xg for 5 min. The plasma was removed, and the cells were washed with PBS, before red blood cells were lysed using ACK buffer (Lonza, Basel, Switzerland) for 4 min and blocked using 10% FBS in PBS. Cells were then stained in PBS using Live-dead fixable Aqua (ThermoFisher Scientific) at 4°C for 15 min. OVA_257-264_ (SIINFEKL)-specific T cells were detected via pentamers staining PE-labelled H2-Kb/OVA_257-264_ (Proimmune, Oxford, UK) in PBS supplemented with 2%, incubated at RT for 30 min. Then cells were stained for surface markers using anti-mouse CD45 (BV421; clone: 30F11), CD3 (BUV396; clone: 17A2; BD Bioscience), CD4 (APC; clone: GK1.5; BioLegend), CD8 (APC-Cy7; clone: 53-6.7), CD44 (PerCP/Cy5.5; clone: IM7), CD62L (FITC; clone: MEL-14) for 15 min at RT at 4°C. Finally, the cells were washed and analysed by flow cytometry using a BD LSRFortessa (BD Biosciences, San Jose, USA). The data were analysed by FlowJo (LLC, Ashland, OR USA).

### Titration of antigen-specific antibodies in the mouse plasma

At day 18 or 55 post-immunisation, 50 μL of mouse blood was collected in EDTA-treated tubes, spun down at 1000 xg for 5 min and the plasma was collected and stored at -80°C until analysis. To measure the antibody titers in the plasma, ELISA plates (Maxisorp, Nunc, Roskilde, Denmark) were coated with 10 μg/mL antigen, either OVA (Sigma-Aldrich, St. Louis, MO, USA) or recombinant VZVO gE (produced as described above) overnight at 4°C. The plates were then blocked using casein (Sigma-Aldrich, St. Louis, MO, USA) for 2 h at RT, after which they were washed with PBS-Tween 0.05% (PBST). The plasma was diluted in casein at a concentration of 1:100 and serially diluted by 10. Samples were applied to the antigen coated plate and incubated at RT for 2 h. The plates were washed three times with PBST and HRP-conjugated antibodies were used to detect antigen-specific antibodies: anti-mouse IgG1 (#1070-05), anti-mouse IgG2a (#1080-05), anti-mouse IgG2b (#1090-05) and anti-mouse IgG3 (#1100-05) from Southern Biotech (Birmingham, AL, USA). Plates were revealed using TMB substrate (EMD Millipore) and stopped with 2N H_2_SO4. We used an Epoch ELISA reader (BioTek, Winooski, VT, USA) to read the absorbance at 450 nm and at 570 nm. The highest plasma dilution for which the corrected absorbance (450 nm-570 nm) was twice the background level defined the antibody titer.

### Restimulation of circulating OVA-specific T cells

Mice were pre-immunised using CpG-adjuvanted OVA as described in “mice immunisation”. At day 35 post-immunisation, about 50 μL of blood per mouse was collected in EDTA-treated tube from the saphenous vein. Red blood cells were lysed using ACK buffer for 3 min, and cells were plated in 50 μL Iscove’s modified Dulbecco’s medium (IMDM) supplemented with 10% FBS and 1% penicillin-streptomycin (P/S) (Gibco, ThermoFisher Scientific, Waltham, MA USA) in a 96 well U-bottom plate. In each well, 50 μL of 1 μg/mL OVA_257-264_ (SIINFEKL) peptide (GenScript) was added using the same medium. The plate was incubated in a cell culture incubator at 37°C, 5% CO_2_ for 4 days. At the end of the culture, the supernatant was collected and IFNγ was quantify using the Mouse IFNγ Quantikine ELISA (R&D systems, Minneapolis, MN USA) according to the manufacturer’s instructions.

### Restimulation of splenocytes using VZVO gE

Mice pre-immunised with Varivax or CpG-B-adjuvanted VZVO gE were boosted with 12 μg gE mixed with 25 μg CpG-B at day 60, and their spleen were collected 5 days later. The splenocytes were obtained by mechanical disruption using a 70 μm cell strainer, and the red blood cells were lysed using ACK buffer for 4 min. Splenocytes were resuspended in IMDM with 10% FBS and 1% P/S at a concentration of 5000 cells/μL, and 100 μL of splenocytes were added per well in a 96 well U-bottom plate. In each well, 100 μL of medium containing 20 μg of gE or no protein (for unstimulated control) was added. Ionomycin/PMA stimulation was used as a positive control. The plate was then placed in a cell culture incubator at 37°C, 5% CO_2_ for 4 days, after what the supernatant was collected. IFNγ was quantified using the Mouse IFNγ Quantikine ELISA (R&D systems).

### Generation of OVA-expressing B16F10 melanoma cells

The different OVA-expressing B16F10 melanoma cells, i.e., B16mOVA^HI/LO^ and B16-OVA^HI/LO^, were generated as described in reference^25^. B16F10 WT parental cells (American Type Culture Collection, Manassas, VA, USA) were genetically modified using lentiviruses with the approval of the biological and safety institutional guidelines of the University of Chicago. For the membrane-bound OVA, the soluble full-length OVA (UniprotKB P01012) was fused to the transmembrane domain of H2-D^B^ at its C terminus. Sequences were cloned in the pLV backbone (Addgene #36804) and lentiviruses were generated by co-transfection with the packaging plasmids pMD2.G (Addgene #12259), pMDLg/pRRE (Addgene #12251) and pRSV-Rev (Addgene #12253), using PEI. Viruses were produced using standard procedures^25,57^ in HEK293-T cells. At the end of the virus production (48-72h after transfection) the medium was collected and filtered at 0.22 μm. After transduction of the B16F10 WT cells with the OVA variants-encoding viruses, B16mOVA^HI/LO^ and B16-OVA^HI^ underwent monoclonal selection using limiting dilution to ensure stable expression over time. The B16-OVA^LO^ cell line was a gift from B. Huard (University of Geneva, Switzerland). All cell lines were cultured in Dulbecco’s Modified Eagle Medium (DMEM) (Gibco, Waltham, MA, USA) with 10% heat-inactivated fetal bovine serum (Gibco) and were tested negative for mycoplasma by PCR.

### Production of recombinant FabTRP-OVA, OVA WT and VZVO-gE

The sequences of TA99 variable regions were also purchased from GenScript and inserted into a plasmid containing the human CH1 (IgG1) and the IgG kappa constant chains. Full-length OVA was subcloned with a C terminal histidine tag, at the C terminus of the human CH1 domain. Both chains were placed under CMV promoters. As a control, full length OVA with a C terminal histidine tag was subcloned in a pXLG house-customed plasmid under the control of a CMV promoter. As to the VZVO gE, the sequence was purchased from GenScript (Piscataway, NJ, USA), and inserted in the pXLG plasmid as well, with a C-terminal histidine-tag. DNA plasmids were amplified by bacterial maxiprep (Nucleobond Xtra Maxi, Macherey-Nagel, Düren, Germany). The plasmids were then transfected in Human Embryonic Kidney (HEK)-293F cells (ThermoFisher Scientific, USA) using polyethyleneimine-mediated transfection (PEI) and cultured for 7 days in FreeStyle 293 medium (ThermoFisher Scientific, USA). The supernatant was then collected and purified using HisTrap 5 mL for the OVA WT and gE, and with an additional purification by HiTrap MabSelect for the FabTRP-OVA (GE Healthcare Life Sciences, USA). Protein purifications were performed using an Akta Pure 25M fast protein liquid chromatography systems (GE Healthcare) according to the manufacturer’s instructions. The purified proteins were dialysed in PBS, filtered sterilised at 0.25 μm, aliquoted and stored at -80°C. Protein purity and size were assessed by SDS-PAGE, and endotoxins level were measured to be < 5 EU/mg using HEK Blue-mTLR4 assays (Invivogen, San Diego, CA, USA).

### OVA expression by quantitative PCR

The mRNA of the different OVA-expressing B16F10 cell lines (about 1-2 million cells) was extracted using the RNeasy total RNA extraction kit (Qiagen, Hilden, Germany). To generate cDNA, reverse transcription was performed using the SuperScript™ IV VILO™ Master Mix (ThermoFisher Scientific). The resulting cDNA was eluted in nuclease-free water to a final concentration of 10 ng/µL and stored at -20°C until future use. Both kits were used following the manufacturer’s instructions. OVA mRNA levels in the different OVA-expressing cell lines were quantified by qPCR using Taqman reactions with the OVAL primer (Gg00366807_m1) and ActB primer (Mm02619580_g1) in the Taqman Universal PCR Master Mix (all from ThermoFisher Scientific). Reactions were performed in a Mastercycler® ep realplex Real-Time PCR System (Eppendorf). Relative expression of the OVA gene in the different B16.OVA variants was calculated using the ΔΔCt method, with Actin B as the housekeeping gene control.

### OVA expression by flow cytometry

OVA-expressing B16F10 cells were plated in a 96-well plate with triplicates at a density of 100,000 cells per well. Then cells were washed twice with PBS. Next, the cells were incubated with 4% paraformaldehyde (PFA) for 10 min, at RT, followed by washing with PBS for 3 times. The cells were incubated with 0.1% Triton X-100 in PBS for 5 min at RT followed by washing with PBS supplemented with 2% fetal bovine serum (FBS) (FACS buffer) for 3 times. The cells then were incubated with rabbit anti-OVA primary antibody (Rb pAb to Ovalbumin, AB181688, Abcam, Cambridge, UK) for 1 h at 4°C. After washing three times with FACS buffer, the cells were incubated with a secondary antibody, anti-rabbit Alexa Fluor 647-conjugated antibody (1:500), for another 30 min under the same conditions. Following three additional washes with FACS buffer, the samples were analyzed using a flow cytometer BD LSRFortessa, and results were analysed with FlowJo.

### Tumour inoculation, treatment, growth and survival measurements

OVA-expressing cell lines were routinely cultured in DMEM + 10% FBS. Cells were detached and rinsed 5 times with PBS in sterile conditions before inoculation in mice. Mice were anesthetized by isoflurane inhalation and injected intradermally with 1.5 million cells, in all experiments in which the effects of the different cell lines were compared (B16mOVA^HI/LO^, B16-OVA^HI/LO^ and B16F10 WT). In experiments in which treatments were assessed, B16F10 WT cells were injected at 250’000 cells. Treatments (i.e., TA99, FabTRP-OVA, IgG2a, OVA, gE, Varivax, PBS as vehicle only) were diluted in PBS to max. 40 μL and administered intratumorally at d2, 4, 6, 8, 10. The proteins were injected at molar equivalent dose of 50 μg of OVA or 63 μg of gE. Varivax was injected at a dose of 13.5 pfu. Anti-TRP1 TA99 (#BE0151) and IgG2a (clone C1.18.4, #BE0085) (InVivoMab, Bio X Cell, Lebanon, NH, USA), was injected at a dose of 100 μg. In some experiments, mice were additionally treated with 200 μg of anti-PD-1 checkpoint blockade (clone 29F.1A12, #BE0273, InVivoMab, Bio X Cell) at day 4 and then repeated every 7 days via peritoneal injections. Tumour growths were then measured every 2 days using a digital caliper, and the volume was calculated as length*width*height*(π/6). Mice were humanely euthanised when turning sick or when the tumour reached 1 cm^3^. The day of euthanasia defined the survival of the mouse, counted as days post-tumour inoculation. Mouse survival was considered as the day of the mouse euthanasia.

### Treatment with depleting antibodies and validation

In some experiments, depleting antibodies were injected intraperitoneally in mice using anti-CD8α (clone 2.43, #BE0061, InVivoMab Bio X Cell) or/and anti-CD4 (clone GK1.5, #BE0003-1, InVivoMab Bio X Cell), at a dose of 500 μg. Injections were performed 3 days before tumour inoculation, and repeated at day 4 and 11 days post-inoculation. To validate proper CD8^+^ and CD4^+^ T cells depletion in mice, blood samples were taken 2 days after the first injection and cells were analysed by flow cytometry according to the procedure described in “Detection of OVA-specific CD8^+^ T cells in the blood”. Cells were stained for Live/dead fixable viability dye Blue (ThermoFisher Scientific) and anti-mouse CD45 (PE; clone: I3/2.3), CD3 (APC; clone: 17A2), CD4 (FITC; clone: GK1.5), and CD8 (APC-Cy7; clone: 53-6.7).

### Immunohistochemistry staining

B16F10 tumours were intradermally injected in mice and collected when the tumour size was about 200-300 mm^3^. The tumours were embedded into a cryomatrix Tissue-Tek OCT (Sakura Finetek, Torrance, CA USA), and rapidly cooled down on dry ice, before being stored at -80°C. Blocks were then sectioned at 6 μm using a cryostat machine (Leica Biosystems, Nussloch, Germany) and placed onto SuperFrost Plus microscope slides (ThermoFisher Scientific). The tumour tissues were then fixed with ice-cold acetone, washed with PBS, blocked with 2% bovine serum albumin (BSA) in PBS for 2 h at room temperature. The slides where then incubated with the anti-TRP1 TA99 antibody (#BE0151) at a final concentration of 1 μg/mL in 0.2% BSA in PBS, or with the control mouse IgG2a antibody (clone C1.18.4, #BE0085) overnight at 4°C. The day after, the slides were washed three times in PBS for 5 min, and incubated 1 h at room temperature with a secondary AlexaFluor 647-conjugated goat anti-mouse. The slides were washed 3 times for 10 min each in PBS and mounted using a fluorescence mounting medium (Dako Omnis; Agilent Technologies, Santa Clara, CA USA) containing 500 ng/mL of DAPI for nuclear staining. The tumours were then imaged with a fluorescent microscope (DMi8, Leica Biosystems) and analysed with Fiji (ImageJ, National Institutes of Health (NIH), Bethesda, MD USA).

### Binding of FabTRP-OVA on B16F10 WT cells

B16F10 WT cells were first stained for live dead in PBS for 15 min on ice. Then, the cells were incubated with 100 nM of FabTRP-OVA or OVA wild-type for 30 min on ice. The cells were then stained using an anti-histidine tag antibody (PE; clone J095G46; BioLegend) to detect the bound proteins, before being analysed by flow cytometry using a BD LSRFortessa. Data were subsequently graphed using FlowJo.

### Analysis of innate immune cells at the tumour site

Skin of mice at the site of tumour injection (about 1 cm^2^) was collected 4 days post-inoculation. The skin was finely minced in small pieces with scissors and transferred into 5 mL tubes containing a magnetic stir bar. Samples were enzymatically digested by collagenase IV (1 mg/mL) and DNAse I (40 μg/mL) in DMEM supplemented with 2% FBS and 1.2 mL CaCl_2_. Digestion was carried out for 1 h at 37°C in a water bath under slow magnetic stirring. The suspension was mechanically dissociated by repeated pipetting (about 100 times). The dissociated cells were then passed through a 70 μm cell strained into a 50 mL tube and washed thoroughly with DMEM with 2% FBS. Undigested skin fragments were subjected to a second enzymatic digestion using collagenase D (3.3 mg/mL) and DNAse I (40 μg/mL) in in DMEM supplemented with 2% FBS and 1.2 mL CaCl_2_. Samples were incubated at 37°C with slow stirring for 15 min, pipetted 100 times, and further incubated for 20 min. EDTA was then added to a final concentration of 5 mM, followed by another round of pipetting. The suspension was filtered through 70 μm strainer and washed, and undigested tissue was discarded. Cells were then pelleted by centrifugation at 400 xg for 5 min, and red blood cells were lysed with 1 mL ACK (Ammonium-Chloride-Potassium) buffer for 3 min. Lysis was quenched with DMEM with 2% FBS, after which the cells were washed, resuspended in 2 mL and filtered again. Cells were then counted and 1 million cells were stained per well. Cells were stained with live dead fixable blue for 15 min in PBS on ice. The cells were then washed and stained for surface markers using anti-mouse CD11b (BV711; clone M1/70; #101241), CD11c (PE-Cy7; clone N418; #117317), CD16/32 (APC-Cy7; clone 93), CD86 (BV421; clone: GL-1), F4/80 (BUV395; clone: T45-2342; BD Bioscience), Ly6C (PE; clone HK1.4), Ly6G (FITC; clone 1A8), I-A/I-E (BV605; clone M5/114.15.2) NK1.1 (PerCP-Cy5.5; clone S17016D) from BioLegend, if not otherwise stated. Cells were then quantified by flow cytometry using a BD LSRFortessa and the data were analysed by FlowJo.

### Data visualisation and statistical analyses

Data visualisation and statistical analyses of the data were performed using Prism 10 (GraphPad, San Diego CA, USA). Outliers were detected using the ROUT tests and excluded from statistical analyses. Used statistical tests and post-tests for multiple comparisons are described in the respective figure legends. Differences between groups were considered statistically significant when the p-value < 0.05. Figures were made using Adobe Illustrator (2023 v27.0, Adobe, San Jose CA, USA).

## REFERENCES

1. Briquez, P. S. et al. Engineering Targeting Materials for Therapeutic Cancer Vaccines. Frontiers Bioeng Biotechnology 8, 19 (2020).

2. Bui, T. A., Mei, H., Sang, R., Ortega, D. G. & Deng, W. Advancements and challenges in developing in vivo CAR T cell therapies for cancer treatment. eBioMedicine 106, 105266 (2024).

3. Zahavi, D. & Weiner, L. Monoclonal Antibodies in Cancer Therapy. Antibodies 9, 34 (2020).

4. Tian, Z., Liu, M., Zhang, Y. & Wang, X. Bispecific T cell engagers: an emerging therapy for management of hematologic malignancies. J Hematol Oncol 14, 75 (2021).

5. Chekaoui, A. et al. Cancer vaccines: an update on recent achievements and prospects for cancer therapy. Clin. Exp. Med. 25, 24 (2024).

6. Samstein, R. M. et al. Tumor mutational load predicts survival after immunotherapy across multiple cancer types. Nat Genet 51, 202–206 (2019).

7. Galluzzi, L., Chan, T. A., Kroemer, G., Wolchok, J. D. & López-Soto, A. The hallmarks of successful anticancer immunotherapy. Sci Transl Med 10, (2018).

8. Vandeborne, L., Pantziarka, P., Nuffel, A. M. T. V. & Bouche, G. Repurposing Infectious Diseases Vaccines Against Cancer. Front. Oncol. 11, 688755 (2021).

9. Baker, R. E. et al. Infectious disease in an era of global change. Nat. Rev. Microbiol. 20, 193–205 (2022).

10. Zuhair, M., et al. Estimation of the worldwide seroprevalence of cytomegalovirus: A systematic review and meta-analysis. Rev. Méd. Virol. 29, e2034 (2019).

11. Millar, D. G. et al. Antibody-mediated delivery of viral epitopes to tumors harnesses CMV-specific T cells for cancer therapy. Nat. Biotechnol. 38, 420–425 (2020).

12. Gastañaduy, P. A. et al. Measles in the 21st Century: Progress Toward Achieving and Sustaining Elimination. J. Infect. Dis. 224, S420–S428 (2021).

13. WHO, W. H. O. Measles cases surge worldwide, infecting 10.3 million people in 2023. https://www.who.int/news/item/14-11-2024-measles-cases-surge-worldwide--infecting-10.3-million-people-in-2023 (2024).

14. Lobo, N. et al. 100 years of Bacillus Calmette–Guérin immunotherapy: from cattle to COVID-19. Nat. Rev. Urol. 18, 611–622 (2021).

15. Redelman-Sidi, G., Glickman, M. S. & Bochner, B. H. The mechanism of action of BCG therapy for bladder cancer—a current perspective. Nat. Rev. Urol. 11, 153–162 (2014).

16. Melero, I. et al. Repurposing infectious disease vaccines for intratumoral immunotherapy. J. Immunother. Cancer 8, e000443 (2020).

17. Newman, J. H. et al. Intratumoral injection of the seasonal flu shot converts immunologically cold tumors to hot and serves as an immunotherapy for cancer. Proc. Natl. Acad. Sci. 117, 1119–1128 (2020).

18. Cao, D. et al. Redirecting anti-Vaccinia virus T cell immunity for cancer treatment by AAV-mediated delivery of the VV B8R gene. Mol. Ther. - Oncolytics 25, 264–275 (2022).

19. Ishida, E. et al. Intratumoral delivery of an HPV vaccine elicits a broad anti-tumor immune response that translates into a potent anti-tumor effect in a preclinical murine HPV model. *Cancer Immunol.*, Immunother. 68, 1273–1286 (2019).

20. Rosato, P. C. et al. Virus-specific memory T cells populate tumors and can be repurposed for tumor immunotherapy. Nat. Commun. 10, 567 (2019).

21. Brown, M. C. et al. Intratumor childhood vaccine-specific CD4+ T-cell recall coordinates antitumor CD8+ T cells and eosinophils. J. Immunother. Cancer 11, e006463 (2023).

22. Wulp, W. van der et al. Antibody-mediated delivery of viral epitopes to redirect EBV-specific CD8+ T-cell immunity towards cancer cells. Cancer Gene Ther. 31, 58–68 (2024).

23. Sethi, S. K. et al. Repurposing anti-viral subunit and mRNA vaccines T cell immunity for intratumoral immunotherapy against solid tumors. npj Vaccines 10, 84 (2025).

24. Tähtinen, S. et al. Exploiting Preexisting Immunity to Enhance Oncolytic Cancer Immunotherapy. Cancer Res. 80, 2575–2585 (2020).

25. Goldberger, Z. et al. Membrane-localized neoantigens predict the efficacy of cancer immunotherapy. Cell Rep. Med. 4, 101145 (2023).

26. Brooks, D. G., Walsh, K. B., Elsaesser, H. & Oldstone, M. B. A. IL-10 directly suppresses CD4 but not CD8 T cell effector and memory responses following acute viral infection. Proc. Natl. Acad. Sci. 107, 3018–3023 (2010).

27. Clynes, R., Takechi, Y., Moroi, Y., Houghton, A. & Ravetch, J. V. Fc receptors are required in passive and active immunity to melanoma. Proc. Natl. Acad. Sci. 95, 652–656 (1998).

28. Pérez-Lorenzo, R., Erjavec, S. O., Christiano, A. M. & Clynes, R. Improved therapeutic efficacy of unmodified anti-tumor antibodies by immune checkpoint blockade and kinase targeted therapy in mouse models of melanoma. Oncotarget 12, 66–80 (2021).

29. They, L. et al. PD-1 blockade at the time of tumor escape potentiates the immune-mediated antitumor effects of a melanoma-targeting monoclonal antibody. OncoImmunology 6, e1353857 (2017).

30. Yang, C. et al. Global, regional, and national burden of varicella-zoster infections in adults aged 70 years and older from 1997 to 2021: Findings from the Global Burden of Disease Study. J. Infect. Public Heal. 18, 102868 (2025).

31. CDC, C. for D. C. and P. Impact of U.S. Chickenpox Vaccination Program. https://www.cdc.gov/chickenpox/vaccination-impact/index.html (2024).

32. Ning, Y. et al. Alphaherpesvirus glycoprotein E: A review of its interactions with other proteins of the virus and its application in vaccinology. Front. Microbiol. 13, 970545 (2022).

33. Zerboni, L., Sen, N., Oliver, S. L. & Arvin, A. M. Molecular mechanisms of varicella zoster virus pathogenesis. Nat. Rev. Microbiol. 12, 197–210 (2014).

34. Hu, W. et al. Redirecting adaptive immunity against foreign antigens to tumors for cancer therapy. Cancer Biol. Ther. 6, 1773–1779 (2007).

35. Çuburu, N., et al. Harnessing anti-cytomegalovirus immunity for local immunotherapy against solid tumors. Proc. Natl. Acad. Sci. 119, e2116738119 (2022).

36. Ostroumov, D., Fekete-Drimusz, N., Saborowski, M., Kühnel, F. & Woller, N. CD4 and CD8 T lymphocyte interplay in controlling tumor growth. Cell. Mol. Life Sci. 75, 689–713 (2018).

37. Narni-Mancinelli, E. et al. Inflammatory Monocytes and Neutrophils Are Licensed to Kill during Memory Responses In Vivo. PLoS Pathog. 7, e1002457 (2011).

38. Haabeth, O. A. W. et al. How Do CD4+ T Cells Detect and Eliminate Tumor Cells That Either Lack or Express MHC Class II Molecules? Front. Immunol. 5, 174 (2014).

39. Bawden, E. G., et al. CD4+ T cell immunity against cutaneous melanoma encompasses multifaceted MHC II–dependent responses. Sci. Immunol. 9, eadi9517 (2024).

40. Akhmetzyanova, I. et al. Tumor-specific CD4+ T cells develop cytotoxic activity and eliminate virus-induced tumor cells in the absence of regulatory T cells. *Cancer Immunol.*, Immunother. 62, 257–271 (2013).

41. Hong, Y. & Kwak, K. Both sides now: evolutionary traits of antigens and B cells in tolerance and activation. Front. Immunol. 15, 1456220 (2024).

42. Snapper, C. M. Distinct Immunologic Properties of Soluble Versus Particulate Antigens. Front. Immunol. 9, 598 (2018).

43. Ackerman, A. L., Kyritsis, C., Tampé, R. & Cresswell, P. Access of soluble antigens to the endoplasmic reticulum can explain cross-presentation by dendritic cells. Nat. Immunol. 6, 107–113 (2005).

44. Biswas, N., Chakrabarti, S., Padul, V., Jones, L. D. & Ashili, S. Designing neoantigen cancer vaccines, trials, and outcomes. Front. Immunol. 14, 1105420 (2023).

45. Kulpa, D. A., Cid, N. D., Peterson, K. A. & Collins, K. L. Adaptor Protein 1 Promotes Cross-Presentation through the Same Tyrosine Signal in Major Histocompatibility Complex Class I as That Targeted by HIV-1. J. Virol. 87, 8085–8098 (2013).

46. Ricca, J. M. et al. Pre-existing Immunity to Oncolytic Virus Potentiates Its Immunotherapeutic Efficacy. Mol. Ther. 26, 1008–1019 (2018).

47. Mok, D. Z. L. & Chan, K. R. The Effects of Pre-Existing Antibodies on Live-Attenuated Viral Vaccines. Viruses 12, 520 (2020).

48. Mohite, P. et al. Revolutionizing Cancer Treatment: Unleashing the Power of Viral Vaccines, Monoclonal Antibodies, and Proteolysis-Targeting Chimeras in the New Era of Immunotherapy. ACS Omega 9, 7277–7295 (2024).

49. Leb-Reichl, V. M. et al. Leveraging immune memory against measles virus as an antitumor strategy in a preclinical model of aggressive squamous cell carcinoma. J. Immunother. Cancer 9, e002170 (2021).

50. Shekarian, T. et al. Repurposing rotavirus vaccines for intratumoral immunotherapy can overcome resistance to immune checkpoint blockade. Sci. Transl. Med. 11, (2019).

51. Aznar, M. A. et al. Repurposing the yellow fever vaccine for intratumoral immunotherapy. EMBO Mol. Med. 12, EMMM201910375 (2019).

52. Zappasodi, R., Merghoub, T. & Wolchok, J. D. Emerging Concepts for Immune Checkpoint Blockade-Based Combination Therapies. Cancer Cell 33, 581–598 (2018).

53. Britsch, I. et al. Novel Fab-peptide-HLA-I fusion proteins for redirecting pre-existing anti-CMV T cell immunity to selectively eliminate carcinoma cells. OncoImmunology 12, 2207868 (2023).

54. Wulp, W. van der et al. Increasing the odds: antibody-mediated delivery of two distinct immunogenic T-cell epitopes with one antibody. OncoImmunology 14, 2508050 (2025).

55. Simoni, Y. et al. Bystander CD8+ T cells are abundant and phenotypically distinct in human tumour infiltrates. Nature 557, 575–579 (2018).

56. Erkes, D. A. et al. Virus-Specific CD8+ T Cells Infiltrate Melanoma Lesions and Retain Function Independently of PD-1 Expression. J. Immunol. 198, 2979–2988 (2017).

57. Brown, L. Y., Dong, W. & Kantor, B. An Improved Protocol for the Production of Lentiviral Vectors. STAR Protoc. 1, 100152 (2020).

