## Supplementary Materials for "Membrane localisation and checkpoint blockade enhance xenoantigen delivery to redirect pre-existing immunity against tumours"

### COMPETING INTERESTS

The authors have no conflict of interest to declare.

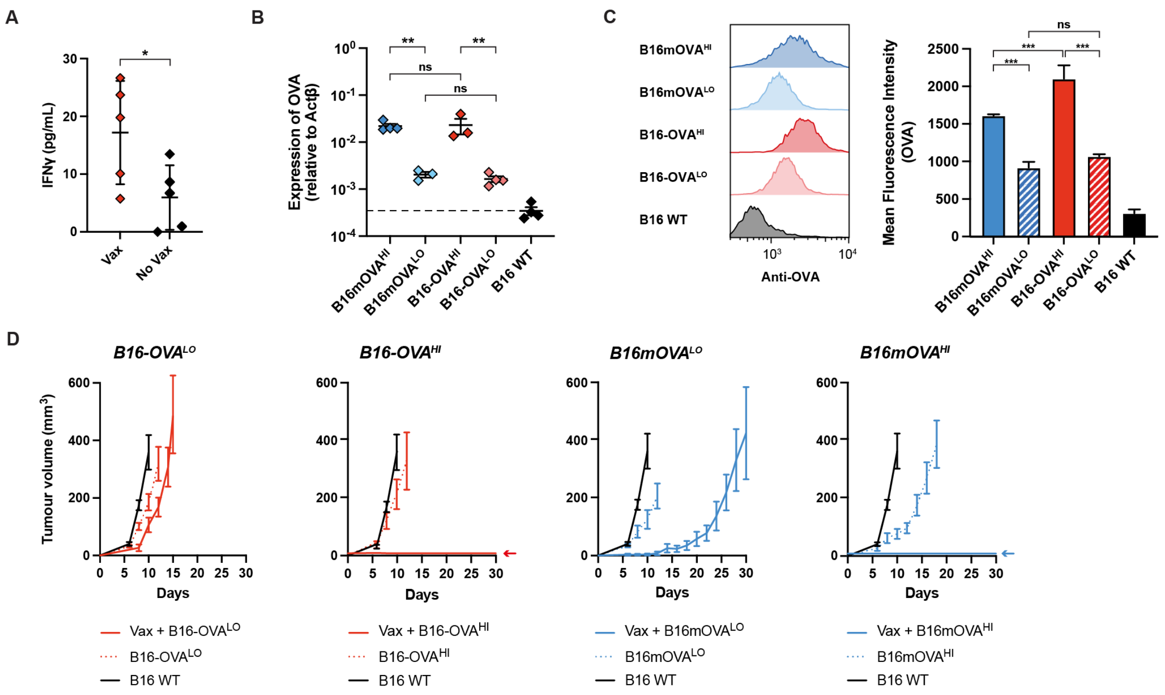


***Figure S1.*** **OVA expression in the engineered B16F10 cell lines and effect of the pre-immunization on OVA-specific T cells and tumour growth.** A) Quantification of IFNγ secreted in the supernatant of T cells collected from the blood (at d35) of pre-immunised (Vax) or naïve (No Vax) mice upon restimulation with $OVA_{257\text{-}264}$ (SIINFEKL) peptide (n = 5, mean ± SD, unpaired t-test). B, C) Quantification of OVA in the OVA-expressing B16F10 cell lines by qPCR (A) and flow cytometry using intracellular detection (B). D) Tumour growth of the OVA-expressing B16F10 inoculated intradermally in pre-immunised (Vax) compared to naïve mice.

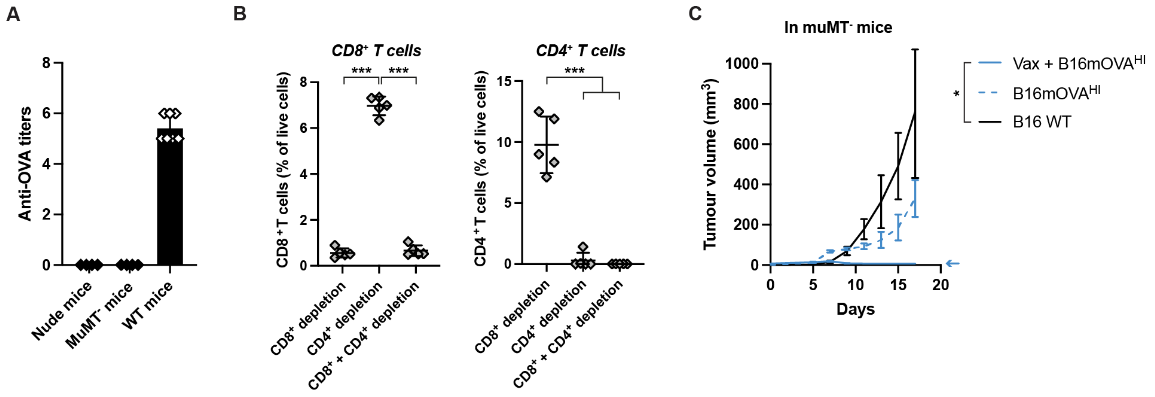


***Figure S2.*** **Depletion of T cells and absence of IgG in B16mOVA$^{HI}$ tumour-bearing mice.** A) Titers of anti-OVA total IgG antibodies at d18 in the nude and MuMT$^{-}$ mice, compared to WT C57BL/6 mice (n ≥ 4, mean ± SD). B) Validation of proper depletion of CD8$^{+}$ T or CD4$^{+}$ T cells in the blood of mice, 2 days after the injection of the depleting antibodies (n = 5, mean ± SD, ANOVA with Tukey's post-test). C) B16mOVA$^{HI}$ growth in pre-immunised or naïve MuMT$^{-}$ mice, which lack mature B cells (n ≥ 3, mean ± SEM, Kruskal-Wallis with Dunn's post-test on d15).

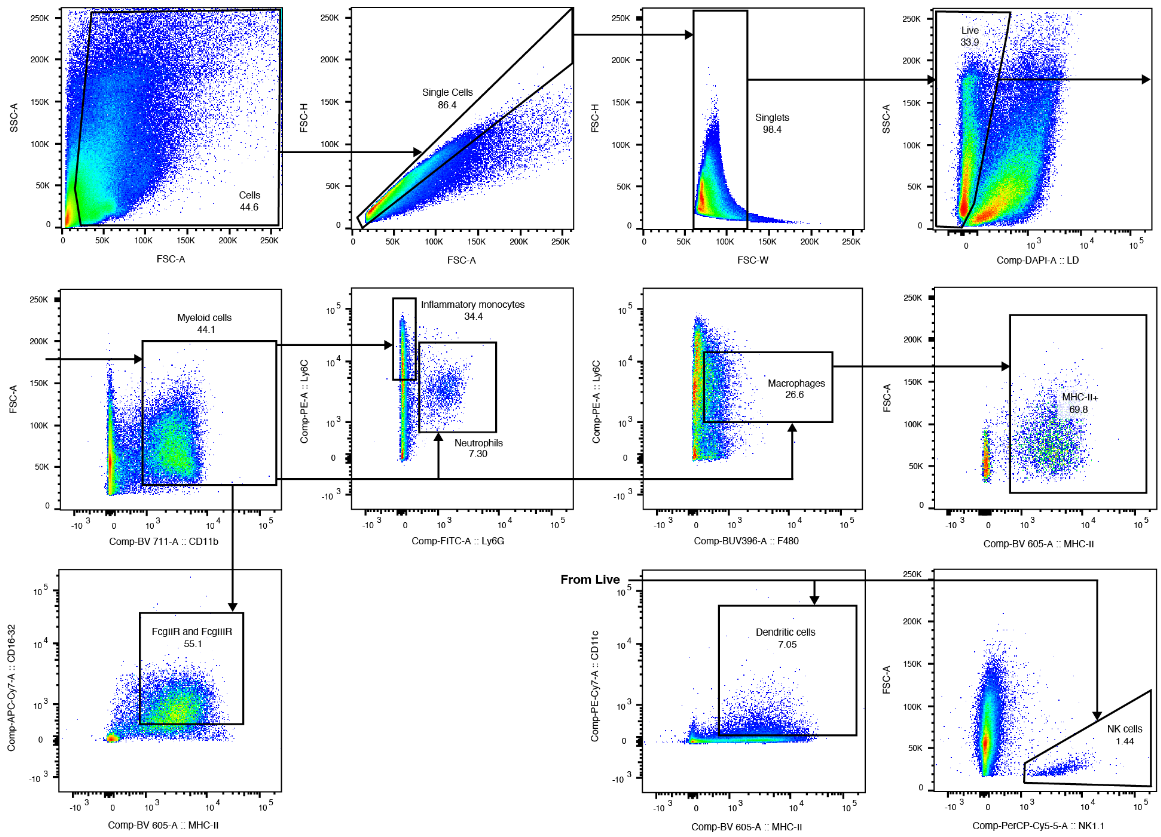


***Figure S3.*** **Gating strategy for the characterization of intradermal innate immune cells at the tumour site 4 days after inoculation.** Flow cytometry was used to quantify the myeloid and NK cells populations 4 days after B16mOVA$^{HI}$ tumour inoculation of in pre-immunised mice. Populations were defined as followed: myeloid cells (CD11b$^{+}$), inflammatory monocytes (CD11b$^{+}$, Ly6C high), neutrophils (CD11b+, Ly6C mid, Ly6G$^{+}$), macrophages (CD11b$^{+}$, Ly6C mid, F4/80$^{+}$), FcγII/IIIR-expressing cells (CD11b+, MHC-II$^{+}$, CD16/32$^{+}$), dendritic cells (CD11c+, MHC-II$^{+}$), NK cells (NK1.1$^{+}$).

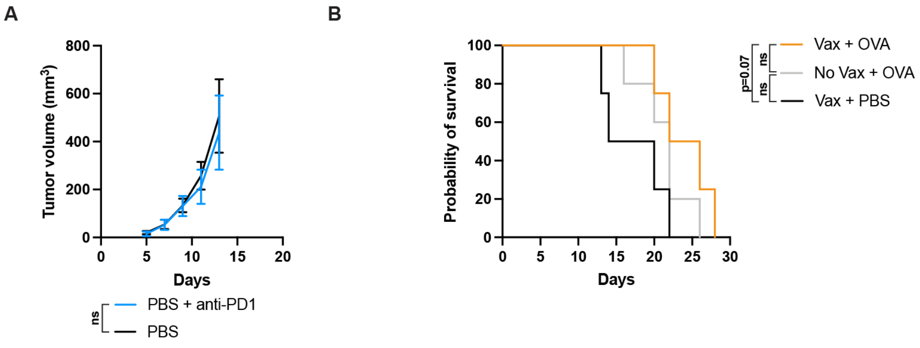


***Figure S4.*** **B16F10 resistance to anti-PD1 therapy and survival of xenoantigen-treated mice**. A) Tumour growth of B16F10 tumours upon treatment with anti-PD-1 therapy (n ≥ 6, unpaired t test on d15). B) Survival of pre-immunised or naïve B16F10 WT tumour-bearing mice treated with OVA or PBS (vehicle only) in combination with anti-PD1 checkpoint blockade (n ≥ 4, log-rank tests).
